# Versatile methanotrophs form an active methane biofilter in the oxycline of a seasonally stratified coastal basin

**DOI:** 10.1101/2022.10.28.513710

**Authors:** Jessica Venetz, Olga M. Żygadłowska, Wytze K. Lenstra, Niels A.G.M. van Helmond, Guylaine H.L. Nuijten, Anna J. Wallenius, Paula Dalcin Martins, Caroline P. Slomp, Mike S.M. Jetten, Annelies J. Veraart

## Abstract

The potential and drivers of microbial methane removal in the water column of seasonally stratified coastal ecosystems and the importance of the methanotrophic community composition for ecosystem functioning are not well explored. Here, we combined depth profiles of oxygen and methane with 16S rRNA gene amplicon sequencing, metagenomics, and methane oxidation rates at discrete depths in a stratified coastal marine system (Lake Grevelingen, The Netherlands). Three amplicon sequence variants (ASVs) belonging to different genera of aerobic *Methylomonadaceae* and the corresponding three methanotrophic metagenome-assembled genomes (MOB-MAGs) were retrieved by 16S rRNA sequencing and metagenomic analysis respectively. The abundances of the different methanotrophic ASVs and MOB-MAGs peaked at different depths along the methane oxygen counter-gradient and the MOB-MAGs show a quite diverse genomic potential regarding oxygen metabolism, partial denitrification, and sulfur metabolism. Moreover, potential aerobic methane oxidation rates indicated high methanotrophic activity throughout the methane oxygen counter-gradient, even at depths with low *in situ* methane or oxygen concentration. This suggests that niche-partitioning with high genomic versatility of the present *Methylomonadaceae* might contribute to the functional resilience of the methanotrophic community and ultimately the efficiency of methane removal in the stratified water column of marine Lake Grevelingen.

## Introduction

Coastal and estuarine-shelf ecosystems contribute up to 75 % of marine methane fluxes to the atmosphere (Upstill-Goddard and Barnes, 2016; Dean *et al*., 2018; Rosentreter *et al*., 2021). Currently, coastal ecosystems are facing warming, enhanced eutrophication, increased stratification and expansion of hypoxic or anoxic zones ((Breitburg *et al*., 2018; Seidel *et al*., 2022). This will likely alter methane cycling and possibly result in larger methane emissions to the atmosphere. The quantification of methane emissions from these ecosystems remains challenging due to the dynamic nature of the occurring biogeochemical processes (Weber *et al*., 2019; Rosentreter *et al*., 2021). Consequently, there is a large uncertainty in estimates of global marine methane emissions (4-27 Tg of CH4 per year) (Weber *et al*., 2019; Saunois *et al*., 2020; Rosentreter *et al*., 2021).

Environmental methane emissions are a result of an imbalance between methane production and methane removal. In coastal ecosystems, methane is produced in the anoxic sediment by methanogenic archaea. Up to 90 % of this methane is estimated to be removed by archaeal methanotrophs in a syntrophic relationship with sulfate-reducing bacteria in the sulfate-methane transition zone (SMTZ) or coupled to various other electron acceptors (Knittel and Boetius, 2009; Wallenius *et al*., 2021). The efficiency of this so-called methane-oxidation filter in the sediment can vary over the season and may lead to high benthic CH_4_ fluxes (Egger *et al*., 2016) and elevated methane concentrations in the water column of hypoxic, stratified coastal waters (Steinle *et al*., 2015; Myllykangas *et al*., 2020; Martins *et al*., 2022). If methane is leaking from the sediment to the bottom water, an active microbial biofilter in the water column can further mitigate methane emissions to the atmosphere, as demonstrated for other coastal ecosystems (Pack *et al*., 2015; Matoušů *et al*., 2017; Steinsdóttir *et al*., 2022).

To improve environmental policy-making and predictions of future methane emissions, the understanding of the microbial methane dynamics in the water column is crucial. The activity and distribution of methanotrophs in the water column are influenced by factors such as oxygen availability, methane concentrations, salinity, temperature, and hydrodynamics (Madigan *et al*., 2006; Steinle *et al*., 2015; Nijman *et al*., 2021). The efficiency and drivers of this methane filter, however, are less well-understood in the water column coastal systems.

Aerobic methanotrophic bacteria (MOB) are phylogenetically distributed among Alphaproteobacteria, Gammaproteobacteria, Verrucomicrobia and the NC10 phylum (Hanson and Hanson, 1996; Op den Camp *et al*., 2009; Ettwig *et al*., 2010; Semrau *et al*., 2010). As a result of the phylogenetic and metabolic diversity (Kits *et al*., 2015; Hisako Hirayama *et al*., 2022; Schmitz *et al*., 2022), different groups of methanotrophs inhabit distinctive niches in the environment. While Verrucomicrobia MOB seem to inhabit extreme environments such as low pH ecosystems (Dunfield *et al*., 2007; Pol *et al*., 2007), alpha-MOBs are mostly found under oligotrophic conditions (Ho *et al*., 2013; Kaupper *et al*., 2020). Gamma-MOB seem to be less resilient to disturbances and are found where methane and other nutrients are not limiting (Ho *et al*., 2013). When methane, oxygen, and nutrient availability in the water column varies with depth during summer stratification, this niche specification within the methanotrophic community may foster efficient methane removal in the water column (Mayr, Zimmermann, Guggenheim, *et al*., 2020). However, we have little insight into the structure of the methanotrophic community in the water column of coastal waters during summer stratification and hypoxia. By unravelling the biogeochemical niches of the present methanotrophic community, we can improve our understanding of what drives methanotrophic activity *in situ* and evaluate the efficiency and stability of the water column microbial methane filter.

Here, we study the role of aerobic methane oxidation in the water column of Marine Lake Grevelingen (NL) during water column stratification in late summer. We present water column profiles of methane and oxygen paired with methane oxidation assays and test the effect of oxygen concentration on methane oxidation potential using batch incubations. In addition, we use 16S rRNA gene sequencing and metagenomics to explore both microbial community composition and the metabolic potential of the methanotrophic communities respectively.

## Results

### Water column chemistry

The methane and oxygen concentrations in the water column of Lake Grevelingen were measured in September 2020. The methane and oxygen profiles showed a clear counter gradient (figure 1A). As the oxygen concentrations remained at around 7 μM below 35 m, we assume that this is rather a common background signal problem of the Seabird oxygen sensor (SBE43) than a real signal. Therefore, oxygen concentrations below 7 μM were considered anoxic. In the anoxic bottom water methane concentrations were high (73 μM). A narrow oxycline was observed between 32 and 35 m with oxygen concentrations from 37.6 μM at 35 m to 115.4 μM at 32 m. Here, the concentration of methane decreased rapidly from 6 μM at 35 m to 0.2 μM at 32 m and remained this high in the surface water.

**Figure 1:**
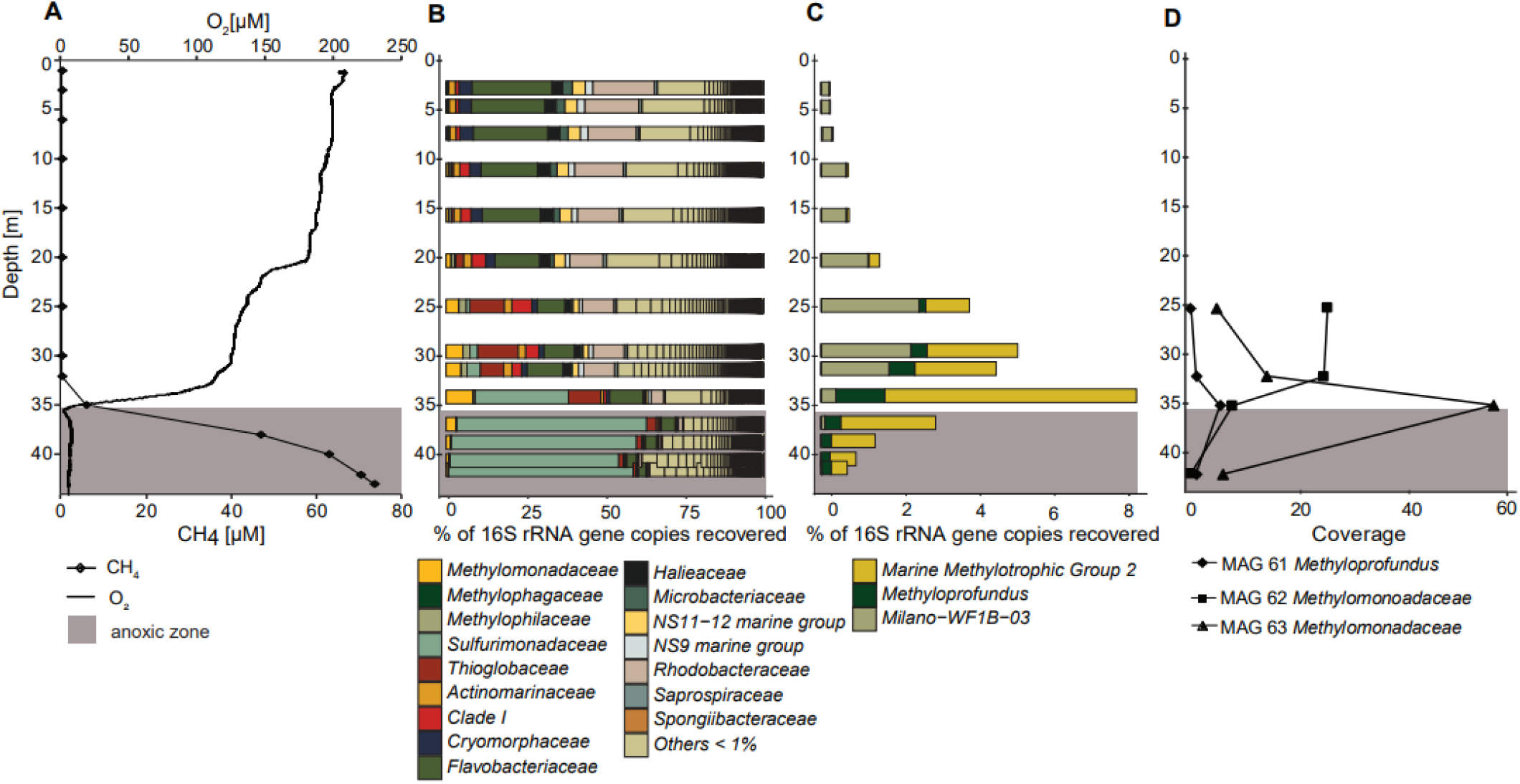
Water column depth profiles of methane and oxygen and the methanotrophic community structure during summer stratification in marine Lake Grevelingen. The grey area indicates anoxic water. **A** methane (diamonds) and oxygen concentrations (black line) with depth in the lake. **B** relative abundance of bacterial families obtained from 16S rRNA sequencing. **C** Vertical distribution of MOB-genera obtained by 16S rRNA sequencing. **D** Vertical distribution of MOB-MAGs obtained from metagenomic analysis.

Concurrent with the oxycline, the temperature decreased at 32 m from 19.9°C in the upper water, to 9.4 °C in the bottom below 38 m (supplemental material, table 1). Salinity showed little change with depth (range of 31 to 32 (supplemental material, table 1). A detailed discussion of the water column chemistry at the time of sampling can be found in Zygadlowska et al., 2022.

**Table 1:** Presence or absence of genes for methane oxidation, oxidases and nitrogen cycling of the different methanotrophic

| Metabolism | MOB-MAG61 | MOB-MAG62 | MOB-MAG63 |
| --- | --- | --- | --- |
| Carbon metabolism |  |  |  |
| Methane monooxidase | pMMO | pMMO | pMMO |
| Methanol dehydrogenase | XoxF/mxaF | XoxF/mxaF | XoxF |
| IV high affinity |  |  |  |
| bd oxidase | 3 of 3 | 0 of 3 | 1 of 3 |
| IV low affinity |  |  |  |
| Cytochrome c oxidase | 3 of 3 | 3 of 3 | 3 of 3 |
| Nitrogen metabolism |  |  |  |
| Nitrite oxidoreductase | non | non | NxrAB |
| Nitrate reductase | non | non | NarGHYZ nxrAB |
| Nitrite reductase | NirB, NirS | NirB | NirB, NirK, NirS |
| Nitric oxide reductase | NorB | non | NorB |
| Nitrogen fixation | NifDH | non | non |
| Sulfur metabolism |  |  |  |
| Sulfur oxidation | soxXYZ, A | soxXYZ, ABCD | soxYZCD |

### Changes in microbial community structure and methanotroph abundance in the stratified water column

The high-resolution depth profile of 16S rRNA gene amplicon sequencing revealed a clear vertical pattern in the microbial community structure of bacteria and archaea in the stratified water column. The aerobic methanotrophic family of *Methylomonadaceae* was found at all water depths. However, the abundance increased from 20 m downwards and peaked at 35 m in the hypoxic zone, reaching an abundance of 8 % of bacterial families (Figure 1B). There, also the anaerobic methane-oxidizing archaeal family of *Methanoperedenaceae* with a relative abundance of 2 % of archaeal families was observed, but they were not found at other depths (supplemental material, figure 1). Although the relative abundance of *Methylomonadaceae* decreased below 35 m, the relative abundance was still considerably high in the anoxic waters with relative abundances of 3 and 1.5 % at 38 and 40 m respectively.

The genus-level vertical distribution of *Methylomonadaceae* revealed three genera, *Methyloprofundus, Marine_Methylotrophic_Group_2* and *Milano-WF1B-03* with a shift in abundance with depth (Figure 1C). *Milano-WF1B-03* was most abundant in the oxic part of the water column, reached a maximum at 25 m and decreased until it disappeared at and below 40 m. *Marine_Methylotrophic_Group_2* and *Methyloprofundus* showed an inverted pattern compared to *Milano-WF1B-03*. Although present in oxic waters, the abundance increased with decreasing oxygen concentrations, peaking at the lower oxycline at 35 m and they remained the dominant methanotrophs in the anoxic bottom water.

### Genomic potential and distribution of methanotrophic MAGs in the oxycline

The genomic metabolic potential of the methanotrophs observed in the stratified water column was elucidated by high throughput shotgun-sequencing of four samples along the oxycline. In line with the 16S rRNA gene sequencing, we retrieved three metagenome-assembled genomes (MAGs) with more than 90 % completeness, that were assigned to the family of *Methylomonadaceae*. As the 16S rRNA ASVs, the abundances of these three methanotrophic MAGs varied along the oxycline (Figure 1 D). The abundance of MOB-MAG61 (representing Methyloprofundus) was the lowest with maximum coverage of 5.7 at 35 m. MOB-MAG62 was most abundant at 25 m with a coverage of 25, although at 35 m the coverage was also high with 24, and close to zero at 42 m. The most abundant MOB-MAG63 peaked at 32 m with a coverage of 54. In line with the suggested niche-partitioning of the methanotrophs, differences in the metabolic potential of each MAG were found (table 1). While all three MOB-MAGs had the particulate methane monooxygenase (pMMO). No soluble methane monooxygenase (sMMO) was found in any of the three MOB-MAGs. The nature of the gene encoding for methanol dehydrogenases (MDH) differed in each MAG. While MOB-MAG61 and MOB-MAG62 encoded for both, La-MDH (*xoxF*) and Ca-MDH (*mxaF*), MAG63 only possessed *xoxF* encoding La-MDH. Cytochrome c oxidase (*caa3*)- the low-affinity oxidase - was present in all MAGs. The high-affinity oxidase bd gene cluster was fully present in MOB-MAG61 and partially in MOB-MAG63. Interestingly, we found genes involved in denitrification in all MOB-MAGs. Moreover, in MOB-MAG63 we found all the genes involved in denitrification except for the nitrous oxide reductase (*nosZ*). MOB-MAG61 was the only one to possess NifH, a diagnostic gene for nitrogen fixation, despite the high ammonium concentration throughout the water column (supplemental material, table 1). All MOB-MAGs showed potential for thiosulfate reduction and sulfide oxidation (*sox*). However, only MOB-MAG62 was equipped with the entire *sox* gene cluster.

### Potential aerobic methane oxidation rates along the methane oxygen counter-gradient

The potential methane oxidation rates were measured at four different depths, corresponding to four different oxygen concentrations (saturated: 270, oxic: 115, hypoxic: 38 and anoxic: <7 μM) and ranged from 0.1 to 4.6 μmol L^-1^d^-1^ (Figure 2). Rates of methane oxidation were highest at 32 m with values ranging from 3.1 to 4.6 μmol L^-1^ d^-1^. Methane oxidation was significantly affected by the interaction between water depth and oxygen (Two-Way ANOVA, F(9, 147) = 2.797, p < 0.005). However, within the treatments for each depth, oxygen concentrations only significantly affected the rates at 35 m (One-way ANOVA, p < 0.0005). In summary, initial *in situ* conditions (e.g. methane and oxygen concentrations) affected methane oxidation rates more strongly than oxygen concentrations in the incubations (see supplemental material, table 2), except for incubations at 5 % O_2_ in the headspace, where methane oxidation rates were not significantly different between depths (p = 0.23).

**Figure 2:**
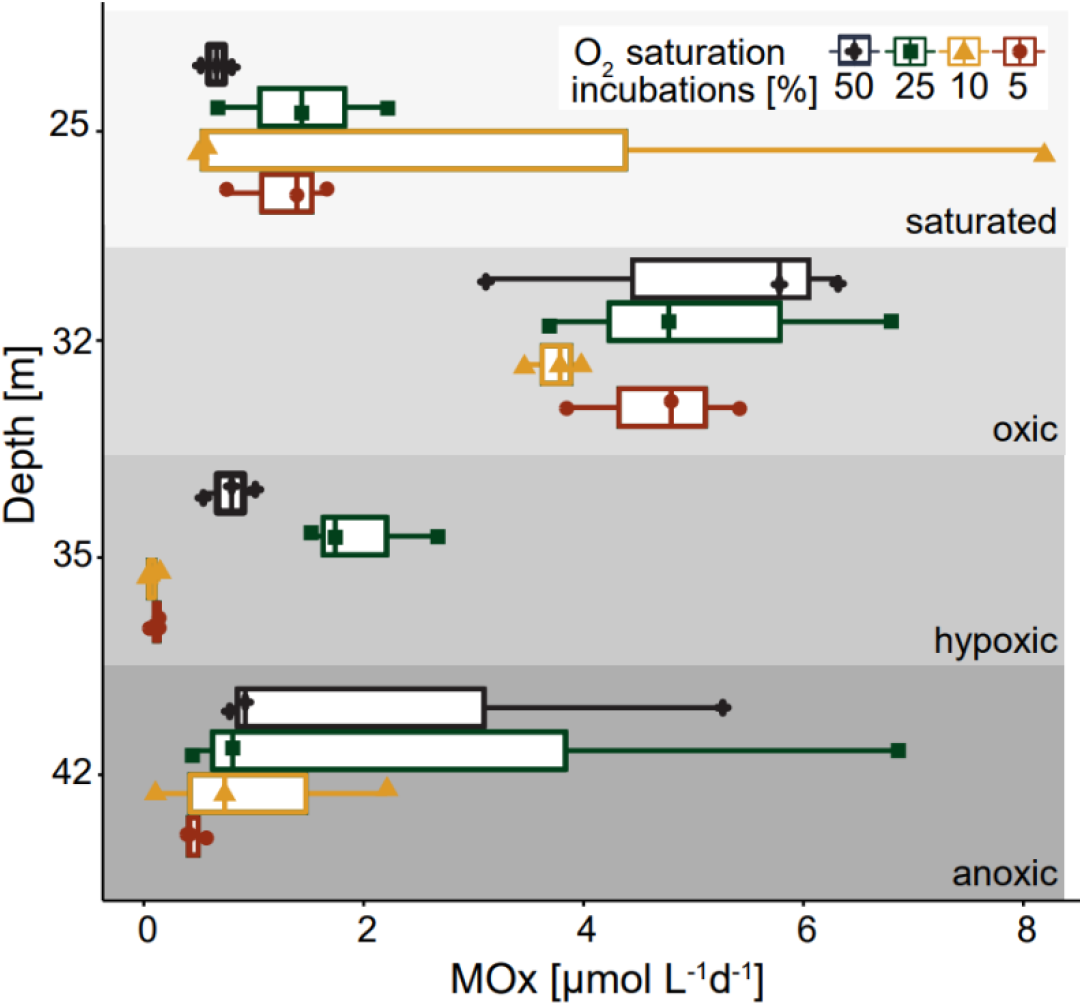
Potential methane oxidation rates [μmol L-1 d-1] of four depths along the oxycline. Each depth was incubated in triplicate with different O_2_ saturations (colors and shapes) and 1 % of ^13^C-CH_4_ to follow ^13^C-CO_2_ production. Grey-scale backgrounds indicate oxygen saturation at the water depth the sample was taken for incubation. Boxes indicate the first and third quartiles, lines indicate the median, whiskers indicate outer data points if less than 1.5*interquartile range from quartiles. Individual data points are shown, n = 48.

## Discussion

### Active microbial biofilter in the oxycline reduces methane emissions

Methane produced in the sediment of coastal ecosystems can subsequently be oxidized anaerobically by archaea with syntrophic sulfate-reducing bacteria in the SMTZ or coupled to various other electron acceptors, reducing methane flux to the overlaying water column. (Knittel and Boetius, 2009; Wallenius *et al*., 2021). In Lake Grevelingen, the bottom water methane concentration was 73 μM, which is up to 100-fold higher than found in comparable ecosystems ((Borges *et al*., 2016; Matoušů *et al*., 2017; Steinsdóttir *et al*., 2022). Here, methane production outpaces methane consumption, likely because of the high sedimentation rate in the Scharendjike basin (∼13 cm yr^-1^ (Egger *et al*., 2016)). Calculated diffusive fluxes of methane from the sediment to the water column range from 0.2–1.0 mol m^-2^ yr^-1^ and the potential for ebullitive flux is high (Egger *et al*., 2016; Żygadłowska *et al*., 2022) which explains these high bottom water methane concentrations. Furthermore, *in situ* methane production by methanogens could potentially add to the methane concentrations, as *Methanosarcinaceae, Methanomicrobiaceae* and *Methanocorpusculaceae* were the dominant archaeal families in the anoxic bottom water (Supplemental material, figure 1).

From this bottom water, methane is transported upwards to the oxycline through vertical mixing, the concentrations decrease moderately. From 35 m upwards, however, the methane concentration decreased strongly from 4 to 0.12 μM at 32 m, resulting in substantially lower methane concentrations above the oxycline. Such sharp methane oxygen counter-gradients in the stratified water column have been found in both marine and freshwater ecosystems, where active microbial methane removal formed a biofilter in the water column (Mau *et al*., 2013; Blees *et al*., 2014; Reis *et al*., 2020).

Methanotrophic *Methylomonadaceae* ASVs were found in the entire water column. Strikingly, the relative abundance peaked concurrently with the strong decrease in methane concentrations within the oxycline. The high relative abundance of aerobic methanotrophic bacteria and the vertical distribution pattern along the methane oxygen counter-gradient in the stratified water column point to an active MOB-biofilter at the oxycline. Nitrate-dependent methanotrophic archaea (*Methanoperedens)* were also present in the hypoxic zone at 35 m, which implies a potential for anaerobic removal of methane. However, we estimate the contribution to methane removal by *Methanoperedens* compared to MOBs to be low, as the abundance of archaea in the water column is 100 times lower than the abundance of bacteria (Thamdrup *et al*., 2019; Steinsdóttir *et al*., 2022). Overall, the methane-oxygen counter-gradient coinciding with the abundance peak of MOB strongly indicates that an active biofilter is effective in removing a large proportion of the high bottom water methane concentrations and thereby lowers total methane emission to the atmosphere.

### Niche partitioning of MOB along the methane-oxygen counter-gradient

In many aquatic ecosystems, the MOB community consists of both gamma and alpha MOB that have different adaptations to different environmental conditions. Gamma-MOB can outcompete alpha-MOB under low methane concentrations (Amaral and Knowles, 1995; Dunfield *et al*., 1999), and are often found in eutrophic aquatic systems (Ho *et al*., 2013). Alpha-MOB seem to be more competitive under oxygen, copper and nitrogen-limiting conditions, e.g. because of their ability to form resting stages and to fix nitrogen (Whittenbury *et al*., 1970; Semrau *et al*., 2010). The methanotrophic community of the water column in Lake Grevelingen mainly consisted of *Methylomonadaceae* belonging to the aerobic gamma-MOB, while the relative abundances of alpha-MOBs never exceeded 0.05 %. Yet, the small reservoir of alpha-MOB indicates that alpha-MOB might still be abundant and active at different stages of water column stratification as observed in freshwater lakes (Mayr, Zimmermann, Dey, *et al*., 2020). Thus, we do not know if gamma-MOB are the dominant methanotrophs in Lake Grevelingen throughout the year or if they were only temporarily dominant during summer stratification.

Gamma-MOB were also the dominant MOB group in the water column of other marine and freshwater ecosystems during water column stratification (Blees *et al*., 2014; Padilla *et al*., 2017; Rissanen *et al*., 2018; Khanongnuch *et al*., 2022). The sole gamma-MOB family detected in the water column of Lake Grevelingen, *Methylomonadaceae*, showed clear niche-partitioning as different genera peaked at various depths. This indicates not only a high ecological versatility among gamma-MOB, which are present at varying conditions such as temperature, pH, high salinity, and low oxygen concentrations (Knief, 2015), but also within the family of *Methylomonadaceae* (Hoefman *et al*., 2014). In eutrophic coastal ecosystems, the metabolic versatility of one methanotrophic family might be especially important for the functional resilience of the ambient methanotrophic community. Assuming that gamma-MOB continuously outcompete other methanotrophs in eutrophic ecosystems, the phylogenetic diversity would be low. This can lower the resilience of the methanotrophic community towards seasonal shifts in e.g. methane and oxygen availability or competition with other members of the microbial community. However, the niche-partitioning of different methanotrophic genera among the *Methylomonadaceae* family indicates a high adaptability of *Methylomonadaceae* to different biogeochemical niches in the water column. This facilitates the formation of a multi-layered, metabolically versatile biofilter and thus enables methane removal across the entire water column. Hence, although summer stratification can lead to the accumulation of methane in the anoxic bottom water, a stable water column stratification may at the same time provide distinct niches with different oxygen and methane availability. Our results show that this possibly fosters pelagic microorganisms to establish a niche-specific population and, in the case of methanotrophs, counteracts high benthic methane fluxes during anoxia.

### Versatile metabolic potential of the methanotrophic community

The potential for niche partitioning of methanotrophs along the methane-oxygen counter gradient of the water column depends on the metabolic versatility of the *in situ* methanotrophic community. In line with our 16S rRNA gene amplicon sequencing results, we retrieved three metagenome-assembled genomes (MAGs) that belonged to the family of *Methylomonadaceae* with abundances peaking at the four sequenced depths along the oxycline. Here we discuss the metabolic potential of the three MOB-MAGs to overcome oxygen limitation through 1) high-affinity oxidases, 2) the use of alternative terminal electron acceptors and/or 3) alternative electron donors.

High- and/or low-affinity oxidases enable the growth of gamma-MOB under varying oxygen concentrations (Oshkin *et al*., 2015). MOB-MAG61 had genes for both, a low-affinity cytochrome c oxidase and cytochrome bd oxidase with presumed high affinity for oxygen. Considering the high relative abundance of MOB-MAG61 in the hypoxic zone at 35 m, it is very likely that the high-affinity oxidase enables these methanotrophs to live off the low amounts of oxygen -supplied through downward turbulent diffusion from the oxygenated water column - as terminal electron acceptor for methane oxidation, even at virtually anoxic conditions. MOB-MAG62 was highly abundant in the oxic water and did not possess this high-affinity bd oxidase but only the low-affinity cytochrome c oxidase. This makes MOB-MAG62 less competitive in the hypoxic zone and is therefore likely outcompeted by MOB-MAG61 and other heterotrophs such as *Flavobacteriaceae* at 35 m. Interestingly, although MOB-MAG63 was most abundant in the hypoxic and anoxic water, only one out of three oxidase-bd subunits were present. Therefore, other metabolic features must explain the high abundance of MOB-MAG63, e.g. the potential to use alternative electron acceptors or alternative metabolic pathways.

Gamma-MOB can oxidize methane under virtually anoxic conditions, (e.g. by using alternative electron acceptors) (van Grinsven *et al*., 2020; Steinsdóttir *et al*., 2022). Some aerobic methanotrophs can use e.g. nitrate as a terminal electron acceptor for the respiratory chain (Kits *et al*., 2015; van Grinsven *et al*., 2020; Steinsdóttir *et al*., 2022). This reduces the need for oxygen substantially as oxygen would only be required for the oxidation of methane to methanol. With nitrate as terminal electron acceptor, a non-oxygen-dependent respiratory mode could be sustained during anoxia (Chen and Strous, 2013), which would explain the potential to oxidize methane in the hypoxic water column (Kits *et al*., 2015; Oswald *et al*., 2016; Rissanen *et al*., 2018). We found all key genes involved in denitrification in MOB-MAG63, except for nitrous oxide reductase. The presence of *narG* and not *napAB* in the MOB-MAG63 further implies that nitrate is potentially used in the respiratory chain, as *nar* contributes to energy conversion and is thus likely involved in energy conservation during oxygen limitation in methanotrophs (Richardson, 2001; Kits *et al*., 2015).

However, as oxygen is still required for the oxidation of methane to methanol, methanotrophs in the anoxic water might make use of alternative electron donors. For instance, methanotrophs may use methanol as an alternative electron donor. While methanol is an important carbon source in the surface ocean (Yang *et al*., 2013), the availability of methanol in the water column is difficult to identify as it is likely rapidly recycled by a variety of microorganisms. We estimate the potential for methanol production below the oxycline of Lake Grevelingen as high. Considering the high sedimentation rate and organic-rich nature of the sediments in this basin (Egger *et al*., 2016), the input of easily degradable organic matter to the anoxic water is expected to be high. Decomposition of this sinking organic material could release methanol (Mincer and Aicher, 2016). MOB-MAG61 and MOB-MAG62 had genes for both the lanthanide and calcium-dependent methanol dehydrogenases (*xoxF* and *maxF*). MOB-MAG63, the most abundant MOB-MAG in the anoxic water, only harboured the lanthanide-dependent methanol dehydrogenase (*xoxF*) which has a higher methanol affinity and yields higher methanol oxidation rates than the calcium-dependent methanol dehydrogenase mxaF (Keltjens JT, 2014). This might enhance the competitiveness to use methanol as an electron donor during hypoxic conditions. As we found the key enzyme sox in all MOB-MAGs, the use of thiosulfate or polysulfide as electron donor is a possibility as was recently shown for a single microorganism (Kojima *et al*., 2014; Gwak *et al*., 2022). However only MOB-MAG62 contained all genes of the *sox* system and peaked in the oxic water column, where thiosulfate would be depleted, and hence thiosulfate is unlikely used as an alternative electron donor in this system. Furthermore, methanotrophs under anoxic conditions could switch to fermentation to maintain a mixture of respiring and fermenting metabolism (Roslev and King, 1994; Kalyuzhnaya *et al*., 2013; Gilman *et al*., 2017).

Despite the metabolic potential and indications for active methane oxidation by *Methylomonadaceae* under anoxic conditions in sediments and the water column of other aquatic ecosystems (Oswald et al., 2016; Steinsdóttir et al., 2022), the stable isotope signatures of the methane profiles in Lake Grevelingen do not indicate high *in situ* activity of anaerobic methane removal in the anoxic bottom water (Żygadłowska et al., 2022). However, the surviving strategies of MOB during, and their ability to re-activate methane oxidation after exposure to anoxia is potentially an important feature to mitigate methane emissions during water column turnover (Blees et al., 2014; Mayr, Zimmermann, Dey, et al., 2020). The activation time and whether the methanotrophs under anoxia remain metabolically active or build resting stages is thus an important topic for future studies.

Overall, the versatile genomic potential and vertical distribution of the three methanotrophic MAGs further confirm the niche-speciation of *Methylomonadaceae* along the methane oxygen counter gradient in the water column of marine Lake Grevelingen. The potential for denitrification implies the capability of the methanotrophic community to oxidize methane under anoxic or hypoxic conditions with alternative electron acceptors. Additionally, the genomic potential to oxidize alternative electron donors further implies potential coping mechanisms during oxygen limitation. We do not know to what extent the ambient methanotrophs in the anoxic zone oxidize methane *in situ* and if methanotrophs oxidize methane with a different electron acceptor or switch to an alternative metabolism entirely. Nonetheless, most importantly, the potential to remain metabolically active during oxygen limitation might ensure a fast re-activation of methane oxidation upon re-oxygenation (e.g. through oxygen intrusion or water column turnover) and therefore might contribute to the functional resilience of the water column methanotrophic community towards oxygen limitation.

### Potential aerobic methane oxidation at different depths and oxygen concentrations

Evidence of active aerobic methane oxidation in the water column of Lake Grevelingen was shown by our oxic incubation experiments. All incubated depths along the methane-oxygen counter gradient oxidized methane at relatively high rates (0.1-4.6 μmol L^-1^ d^-1^). The methane oxidation rates at the different depths were significantly different from each other. The highest methane oxidation rates in the water column of Lake Grevelingen were observed at 32 m where oxygen and methane were readily available, potentially creating an ideal niche for methanotrophs. This is in line with other studies that found the highest rates of methane oxidation in the upper part of the methane oxygen counter-gradient (Cabrol et al., 2020). Potential methane oxidation rates in the hypoxic lower part of the oxycline were lower, despite the higher relative abundance of methanotrophic ASVs at this point. The methane oxidation rates in the anoxic zone at 42 m indicate that methane oxidation can be activated by oxygen within 24 hours, even in an anoxic water column that has been anoxic for weeks to months. Vice-versa, methane oxidation in the oxygenated water at 25 m could be activated by methane addition as well, although the methane profile did not indicate active methane removal at this depth. This suggests that methanotrophs throughout the water column of Lake Grevelingen can quickly adapt to changes in methane and oxygen availability as observed in other ecosystems (Blees et al., 2014; Mayr, Zimmermann, Dey, et al., 2020). Together with the molecular biological data and the chemical profiles, the incubations confirm the high potential of the methane oxidation filter in the water column.

The drivers for *in situ* methane oxidation, and how the methanotrophic community structure and potential methane oxidation rates shift during the seasons, still require further investigation.

Oxygen and methane are presumably the main drivers for methane oxidation in aquatic ecosystems, which is in line with the measured differences in methane oxidation rates between the incubated depths along the methane oxygen counter-gradient. However, varying initial oxygen saturations of 5, 10, 25 and 50 % (corresponding to 11, 23, 57 and 114 μM [O_2_]_liq_) in our incubations, did not influence methane oxidation rates significantly at most depths.

Although this does not exclude oxygen availability as a driver for methane oxidation *in situ*, it improves our understanding of to what extent the ambient methanotrophic community can adapt to changes to *in situ* oxygen supply. Certainly, the relatively high oxygen concentrations in our incubations do not elucidate the effect of oxygen concentrations on enzyme kinetics of methane oxidation, but it sheds light on the potential of methanotrophs in the water column to compete for oxygen with heterotrophs, especially at low oxygen concentrations. At 35 m we observed significant effects of oxygen availability on methane oxidation rates. There, incubations with 5 and 10 % oxygen saturation (11 and 23 μM [O_2_]_liq._), resulted in lower methane oxidation rates compared to incubations with 25 and 50 % (57 and 114 μM [O_2_]_liq._). The dominant methanotroph at 35 m was MOB-MAG63, which showed high genomic potential for methane oxidation with nitrate. The *in situ* oxygen concentrations at this depth (37 μM) might be high enough to activate methane oxidation to methanol, and methanol could further be oxidized with nitrate. Yet, lower oxygen concentrations in our incubations (11 and 23 μM), resulted in significantly lower methane oxidation rates than with 57 and 114 μM. The overarching regulator for this could be the competition for oxygen with heterotrophs, which may be more efficient in scavenging any available oxygen compared to methanotrophs. MOB-MAG63 for instance did not harbour the high-affinity oxidase, which reduces the competitiveness for oxygen scavenging. Under high competitive pressure, it may be energetically better to remain in the anaerobic metabolism until a threshold oxygen concentration induces the transcription of enzymes involved in oxic methane oxidation. Methanotrophs with the genomic potential of MOB-MAG63, could for instance oxidize methanol or sulfur compounds with nitrate under these low oxygen concentrations or revert to a partial fermentative metabolism, to endure oxygen limitation. The low initial oxygen concentrations in our incubations might not have reached the threshold to re-activate aerobic methane oxidation, as oxygen consumption of other microbes was potentially high and decreased the total available oxygen. Distinctively, in the incubations with 50 and 25 % oxygen saturation (114 and 57 μM [O_2_]_liq._), total oxygen concentrations could have remained high enough to reach this threshold, even with cooccurring heterotrophic oxygen consumption, which resulted in higher methane oxidation rates. Intriguingly, methane oxidation rates with 50 % initial oxygen saturation were lower than at 25 %. This non-linear correlation between oxygen concentrations and methane oxidation rates has been shown in other ecosystems (Thottathil *et al*., 2019), which indicates limitations of another compound or inhibition. At 42 m where methanotrophs had been exposed to anoxia for months, higher oxygen concentrations resulted in slightly higher methane oxidation rates, but the differences were not significant. As heterotrophs are less abundant in the anoxic water than at the oxic-anoxic interface, methanotrophs in the anoxic water are less likely outcompeted by active heterotrophs upon re-oxygenation. Thus, the potential to use alternative electron acceptors for methane oxidation of the bottom water methanotrophic community might be less important for efficient methane removal than the ability to enter a resting state and re-activate methane oxidation upon re-oxygenation. In that case, the effect of oxygen concentrations on methane oxidation rates is influenced by the competition for oxygen with other microorganisms and the active metabolic state of the methanotrophs *in situ*. This highlights the importance of measuring methane oxidation rates at a high vertical resolution in the stratified water column and taking the entire microbial community into account, to improve estimations of how much methane is removed and what factors are driving methane oxidation.

## Conclusion

We showed that an active microbial methane filter in the water column of marine Lake Grevelingen, counteracts high benthic methane fluxes during summer stratification and hypoxia. Illumina 16S rRNA gene sequencing analysis showed that *Methylomonadaceae* were abundant in the entire water column with a striking distinct abundance peak coinciding with a sharp methane oxygen counter-gradient. Although the only methanotrophic family, different genera of *Methylomonadaceae* inhabited the potential ecological niches within the stratified water column. Furthermore, three MOB-MAGs belonging to *Methylomonadaceae*, with versatile metabolic potentials and vertical distribution were retrieved. The presence of high-affinity oxidase and genes involved in nitrate reduction and sulfur oxidation in the three MOB-MAGs shed light on potential strategies to overcome temporary oxygen limitation. In fact, potential methane oxidation rates were high in all incubated depths, even in the anoxic bottom water. Moreover, although potential aerobic methane oxidation rates differed at different depths within the methane oxygen counter-gradient, incubations of each depth with various oxygen concentrations did not show any influence of oxygen availability on methane oxidation at most depths.

With these results we conclude that the versatile metabolic potential of members of *Methylomonadaceae* in the stratified water column of Lake Grevelingen 1) enables the population of different geochemical niches within the water column, 2) broadens the vertical range of the active methane oxidation filter, 3) improves the competitiveness, and 4) increases the resilience of the *in situ* methanotrophic community towards changes in the oxygen availability. Thus, the metabolic versatility of the ambient methanotrophic community is a crucial factor for the mitigation of methane emissions from seasonally stratified coastal ecosystems and needs further attention. The investigation of the seasonal dynamics of methane oxidation including the metabolic potential of the methanotrophic community will further improve our understanding of the functioning and drivers of the microbial methane filter in the water column of these highly dynamic coastal ecosystems.

## Experimental procedures

### Fieldwork location and sampling methodologies

Lake Grevelingen is a former estuary in the southwest Netherlands. The lake has a surface area of 115 km^2^ with an average water depth of 5.1 m. Here, we sampled Scharendijke basin, which is part of a former tidal channel and has a depth of 45 m. A more detailed description of the system and the study site can be found elsewhere (Egger et al., 2016; Żygadłowska et al., 2022). Lake Grevelingen was chosen to investigate microbial methane cycling as previous studies indicated that methanogenesis exceeded methane oxidation in the sediment during summer stratification, resulting in a periodic release of methane into the water column (Egger et al., 2016).

We took water column samples at the deepest point in Scharendijke basin (51.742°N; 3.849°E, 45 mbs) (Figure 3), in September 2020 during a two-day sampling campaign with RV Navicula. To determine the extent of stratification of the water column in September 2020 we measured salinity, temperature, and depth with a CTD unit (SBE 911 plus, Sea-Bird Electronics Bellevue (WA), USA). The oxygen distribution was simultaneously recorded by a seabird sensor. Water samples were taken at 14 depths with an 8 L Niskin bottle. The distribution of the sampling depths is depicted in supplement S1.

**Figure 3:**
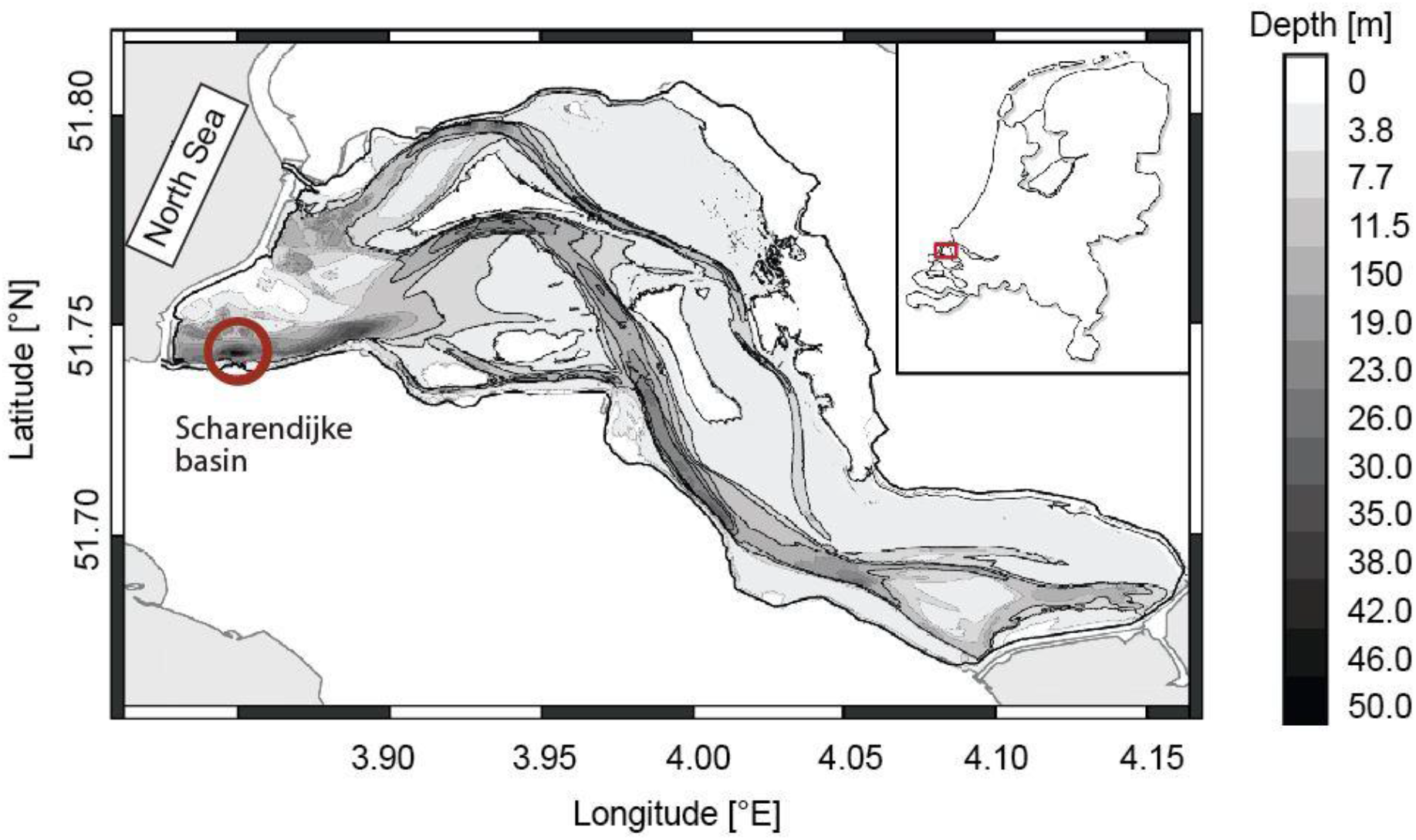
Map of Marine Lake Grevelingen modified from Egger et al., 2016. Sample location Scharendijke (circle) (51.742N;3.849E)

For DNA samples, we filled two sterile 1 L plastic bottles with unfiltered water from each depth and stored them at 4 °C until further processing (within 24 hours).

Samples for methane concentrations were taken by filling 120 ml borosilicate serum bottles (duplicates) directly via gas-tight tubing. To prevent air contamination, the bottle was filled up from the bottom while letting the water overflow three times. Then the bottle was crimp-capped with a butyl stopper and aluminium cap and stored upside down to avoid air contamination during storage. Subsequently, 1 ml of anoxic 50 % ZnCl_2_ solution (w:v) was added to stop microbial activity, and samples were stored at room temperature until further processing.

To test the activity and the response to different oxygen concentrations of the MOB community, we took four additional samples along the oxycline at 20, 32, 35, and 42 m depth. Samples were filled into sterile 1 L borosilicate Schott bottles and stored at 4 °C in the dark.

### Methane concentration measurements

To determine methane concentrations in the water column, 5 ml of nitrogen gas was added to all the methane samples simultaneously removing the same volume of the sample. After equilibrating for at least 2 h methane concentrations were measured with a Thermo Finnigan Trace™ gas chromatograph (Flame Ionization Detector).

### DNA extraction, 16S rRNA gene sequencing, and data analysis

DNA from the water column samples was obtained by filtering between 0.5 to 1 L of water on Supor® PES 0.22 μm filters using a vacuum pump, within a day after sampling. The filters were then stored at -80 °C until further use. DNA was extracted following the protocol of the DNeasy Power water DNA Isolation kit (Qiagen, Venlo, the Netherlands).

The composition of the microbial community was analyzed, by sequencing the V3-V4 region of the 16S rRNA gene (Illumina MiSeq platform, Macrogen, Amsterdam, the Netherlands), using the primer pairs Bac341F (CCTACGGGNGGCWGCAG) (Herlemann et al. 2011) with Bac806R (GGACTACHVGGGTWTCTAAT) (Caporaso et al. 2012) for bacteria, and Arch349F (GYGCASCAGKCGMGAAW) (Takai Ken and Horikoshi Koki, 2000) with Arch806R (GGACTACVSGGGTATCTAAT) (Takai Ken and Horikoshi Koki, 2000) for archaea. RStidio was used to process 16S rRNA sequencing data. First, primers were removed with cutadapt (Martin and Rahmann, 2012) using the options -g, -G, and --discard-untrimmed. Reads were truncated to a length of nt 270 forward nt 260 reverse by following the DADA2 pipeline (Callahan *et al*., 2017) and low-quality reads were removed. Sequences were dereplicated after error 252 models were built. Amplicon sequence variants (ASVs) were then inferred and forward and reverse reads were merged and chimaeras removed. the 254 Silva non-redundant train set v138 downloaded from https://zenodo.org/record/3731176#.XoV8D4gzZaQ was used for taxonomic assignment. Phyloseq was then used for further clustering and calculation of relative abundance. Plots were made with ggplot2.

### Metagenome sequencing, genome binning, and sequence analysis

For deeper insight into the metabolic potential of MOBs across the oxycline in the water column of Lake Grevelingen, samples from 25, 32, 35 and 42 m (congruent with the samples used for the incubations) were sent for full metagenome sequencing (TruSeq Nano DNA (350)kit, Novaseq platform, Macrogen, Amsterdam, the Netherlands). The metagenomic data were analyzed with an in-house automated pipeline “BinMate”. FASTQC was used to assess the quality of the reads and trimming of adapters and low-quality bases were performed with BBTools. Reads were assembled into contigs with megahit. The coverage of each contig was obtained by mapping the reads back to contigs with bbmap. For optimized binning results, BinMate used CONCOCT, MaxBin2 and MetaBAT to perform metagenomic binning. Bins were then dereplicated with DAStool. Final quality checks were performed with CheckM. Medium quality was defined as > 50 % completeness and <10 % contamination. Bins were classified taxonomically with gtdb-tk and annotated with DRAM.

### Potential methane oxidation rates and oxygen manipulation experiments

To investigate the potential activity of methanotrophs and the effect of oxygen on the efficiency of methane removal in the water column of Lake Grevelingen, we conducted methane oxidation experiments in batch incubations. Within 24 h after sampling, 20 ml of each water sample was transferred into autoclaved 120 ml borosilicate serum bottles, closed with Bromo-butyl stoppers, and crimped with an aluminium cap. To remove the ambient gases, the bottles were purged with argon gas for 15 min.

Each of the 4 depths was provided with four different oxygen saturations of 50, 25, 10 and 5 % and 1 % headspace ^13^C-CH_4_ resulting in a dissolved oxygen concentration of 114, 57, 23 and 11 μM and methane concentration of 8 μM respectively. All bottles were incubated under constant shaking (150 rpm) at room temperature in the dark for 95 h.

To follow the oxidation of labelled methane, the increase of produced ^13^C-CO_2_ was measured directly from the headspace with a GC-MS (Agilent 5975C inert MSD). In addition, we measured methane headspace concentrations with a GC (). The total concentrations of ^13^C-CO_2_ and ^13^C-CH_4_ in the incubation bottles were calculated using Henry’s law (see supplements). The rates were then calculated using linear regression of the first linear increase in ^13^C-CO_2_ after the lag phase. To correct for ^13^C-CO_2_ production from other processes such as respiration, non-methanotrophic ^13^C-CO_2_ was calculated as 1.2 % of produced total CO_2_ and subtracted from total ^13^C-CO_2_ production (supplemental material, section 3).

To observe potential changes in the microbial community during the incubations, the triplicate samples were pooled and filtered with a 0.2 μm Supor® PES filter with a vacuum pump and stored at -80 °C until DNA extraction and subsequent sequencing of 16S rRNA gene.

### Statistical analysis

Statistical analyses to test the effect of sampling depth and initial headspace oxygen concentrations on methane oxidation rates were performed in Rstudio with the following packages tidyverse, ggpubr and rstatix.

We used mixed two-way ANOVA followed by post hoc tests. The assumption of normal distribution was tested with the Shapiro-Wilk normality test after log transformation. All data points were normally distributed (p>0.05) except for 35 m 1 % (p = 0.0139). However, we did not find any extreme outliers using the identify_outliers function. Therefore, we decided to not exclude any rates from the biological replicates and considered them as true variations. The homogeneity of variance was tested with Levene’s test and no statistically significant differences were found (p > 0.05).

For the two-way mixed ANOVA test, we used the anova.test() function (depth=within-subject variable, oxygen concentrations = between-subject variable). After a significant two-way interaction was found (p<0.05), we performed a simple main effect analysis and a simple pairwise comparison as post hoc tests.

## Acknowledgements

We thank the crew from the RV Navicula for their enthusiastic cooperation and help. Further, we want to thank Theo Van Alen for the assistance with obtaining 16S rRNA and metagenomic sequencing data.

This study was funded by NESCC NWO/OCW 024002001, ERC Synergy MARIX 854088 and SIAM NWO/OCW 02002002.

