## Supplementary material for "Versatile methanotrophs form an active methane biofilter in the oxycline of a seasonally stratified coastal basin": Suplemental material

### Supplements

#### 1. Water column profile

**Table 1:** *Concentrations of oxygen, methane, nitrate, nitrite and ammonium and Temperature and Salinity at the sampled depths.*

| depth [m] | $O_2$ [µmol $L^{-1}$] | $CH_4$ [µmol $L^{-1}$] | $NO_3^-$ [µmol $L^{-1}$] | $NO_2^-$ [µmol $L^{-1}$] | $NH_4^+$ [µmol $L^{-1}$] | Temp. [°C] | Salinity |
|---|---|---|---|---|---|---|---|
| 1 | 209,0 | 0,2 | 2,0 | 0,7 | 31,4 | 19,7 | 31,5 |
| 3 | 208,4 | 0,2 | 1,9 | 0,6 | 24,3 | 19,7 | 31,5 |
| 6 | 201,0 | 0,2 | 1,7 | 0,6 | 23,3 | 19,7 | 31,5 |
| 10 | 198,9 | 0,2 | 1,8 | 0,5 | 25,4 | 19,7 | 31,6 |
| 15 | 189,7 | 0,3 | 1,5 | 0,5 | 25,9 | 19,7 | 31,6 |
| 20 | 171,3 | 0,2 | 0,6 | 0,4 | 30,1 | 20,0 | 31,9 |
| 25 | 134,6 | 0,1 | 0,5 | 0,3 | 45,4 | 20,1 | 32,0 |
| 30 | 125,2 | 0,1 | 1,9 | 0,5 | 42,2 | 20,0 | 32,1 |
| 32 | 115,4 | 0,2 | 0,8 | 0,5 | 50,0 | 19,9 | 32,1 |
| 35 | 37,6 | 6,0 | 1,8 | 0,5 | 64,0 | 17,9 | 32,0 |
| 38 | 7,3 | 47,0 | 0,4 | 0,3 | 158,6 | 10,8 | 31,4 |
| 40 | 6,8 | 63,0 | 0,3 | 0,4 | 203,9 | 9,6 | 31,3 |
| 42 | 6,3 | 70,3 | 0,3 | 0,3 | 215,9 | 9,3 | 31,2 |
| 43 | 6,2 | 73,5 | 0,9 | 0,4 | 226,5 | 9,1 | 31,2 |

#### 2. 16S rRNA gene and metagenomic sequencing

##### 2.1. Archaeal 16S rRNA gene abundance

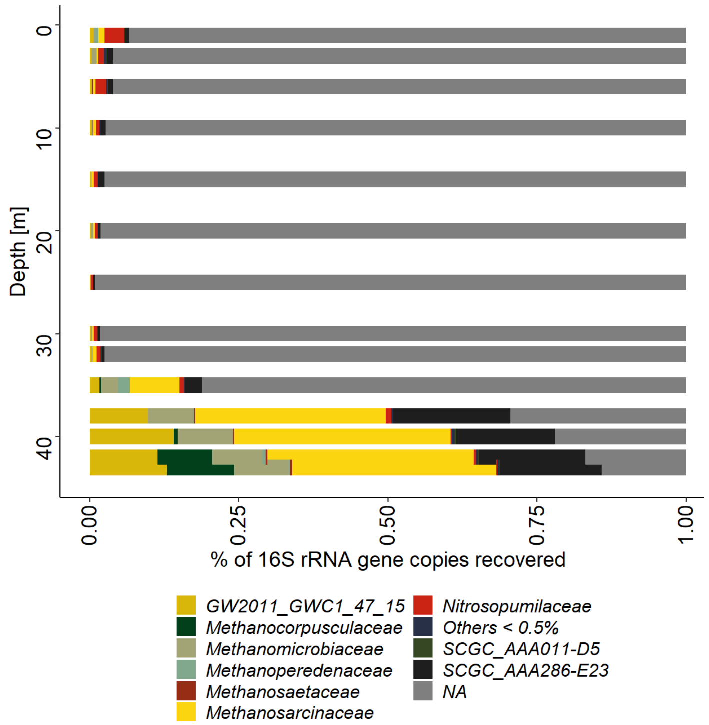


**Figure 1:** *Distribution of archaeal families along the water column profile.*

#### 3. Incubations

##### 3.1. Calculations of gas concentrations

From headspace measurements, we calculated the total gas concentration in the incubations with Henry's law. The concentration of a gas in the aqueous phase ($c_a$) in a closed system can be calculated by multiplying the concentration of the gas phase ($c_g$) with the Henry solubility coefficient $H^{cc}$ (Sander, 2015):

$$c_a H^{cc} c_g$$

The Henry solubility coefficient is defined as follows: $H^{cc} = H^{cp} RT = \beta \frac{1}{RT^{STP}} RT$

$H^{cp}$: Henry solubility coefficient (defined as $c_a$/p)

R: ideal gas constant (8.314 J $mol^{-1}$ $K^{-1}$)

T: temperature (294.14 K)

$T^{STP}$: the standard temperature for Bunsen coefficient (273.15 K)

$\beta$: Bunsen coefficient (including salinity and temperature)

To include the salinity and temperature in the solubility coefficient we calculated the salinity and temperature-dependent Bunsen coefficients with the following equation (Weiss 1970):

$$ln\beta = A_1 + A_2\left(\frac{100}{T}\right) + A_3 ln\left(\frac{T}{100}\right) + S\left[B_1 + B_2\left(\frac{T}{100}\right) + B_3\left(\frac{T}{100}\right)^2\right]$$

$A_{1\text{-}3}$, $B_{1\text{-}3}$: Bunsen constants, specific for gas

T: Temperature (294.15 K)

S: Salinity (30 ‰)

The Bunsen coefficient for $CH_4$, $CO_2$ and $O_2$ was calculated according to the specific constants for each gas (Weiss, 1970, 1974; Yamamoto *et al.*, 1976).

##### 3.2. Correction for naturally produced $^{13}C\text{-}CO_2$

As we could not exclude concomitant respiration in our incubations, we corrected for potentially produced $^{13}C\text{-}CO_2$ by respiration. Therefore, we normalized the total produced $^{13}C\text{-}CO_2$ to the baseline ratio of $^{13}C\text{-}CO_2/^{12}C\text{-}CO_2$ of 1.2 % with the following calculation:

$$^{13}CO_2CH_4oxidation = {}^{13}CO_{2total} - {}^{12}C\frac{O_2 * 1.2\%}{98.8\%}$$

##### 3.3.Methane oxidation rates

**Table2:** *Methane oxidation rates (MOx) together with in situ oxygen saturation of the sampled depths and oxygen saturation in*

| Depth [m] | $O_{2\ in\ situ.}$ [%] | $O_{2\ inc.}$ [%] | MOx [µmol $L^{-1}$ $d^{-1}$] |
|---|---|---|---|
| 25 | 57 | 5 | 1.1 ± 0.3 |
| | | 10 | 2.8 ± 3.2 |
| | | 25 | 1.2 ± 0.5 |
| | | 50 | 0.5 ± 0.1 |
| 32 | 46 | 5 | 4.0 ± 0.6 |
| | | 10 | 3.1 ± 0.2 |
| | | 25 | 4.5 ± 1.0 |
| | | 50 | 4.6 ± 1.1 |
| 35 | 12 | 5 | 0.1 ± 00 |
| | | 10 | 0.1 ± 0.0 |
| | | 25 | 1.6 ± 0.4 |
| | | 50 | 0.7 ± 0.2 |
| 42 | 2 | 5 | 0.4 ± 0.1 |
| | | 10 | 0.8 ± 0.8 |
| | | 25 | 2.3 ±2.6 |
| | | 50 | 1.9 ± 1.8 |

##### 3.4.Statistics

The pairwise T-test between treatments revealed that methane oxidation rates of incubations with 1 % $O_{2\ headspace}$ were significantly different from incubations with 5 % and 10 %, (p= ..) as well as methane oxidation rates of incubations with 2 % $O_{2\ headspace}$ were significantly different from incubations with 5 % and 10 %. Also, 5 % and 10 % were significantly different, but the test showed no statistical difference between methane oxidation rates at 1 % and 2 % headspace oxygen.

Sampling depth, on the other hand, had more influence on methane oxidation rates. The methane oxidation rates of each initial oxygen headspace concentration were strongly and significantly different between all depths, except for 5 % $O_{2\ headspace}$. For incubations with 5 % $O_{2\ headspace}$, methane oxidation rates were not significantly different between the depths (p =0.23). The pairwise T-test, however, revealed that in incubations with 1 % $O_{2\ headspace}$, methane oxidation rates were strongly influenced by depth. In incubations with 2 % $O_{2\ headspace}$, only methane oxidation rates at 35 m were different from 25 and 32 m. At 5 % $O_{2\ headspace}$, all were significantly different and a 10 % only 32 m was different from 25, 35 and 42 m.
